## supplemental Files for "Asymmetric conformations and lipid interactions shape the ATP-coupled cycle of a heterodimeric ABC transporter"

### Supplementary Table 1: Cryo-EM data collection, refinement, and validation statistics

| **Data collection and processing** | | | | | | | |
|---|---|---|---|---|---|---|---|
| Dataset | Dataset 1 | Dataset 2 | Dataset 3 | Dataset 4 | | Dataset 5 | |
| Construct | BmrCD*-QQ | BmrCD*-QQ | BmrCD*-QQ | BmrCD*-QQ | | BmrCD-WT | |
| Lipids used | PC/PA | PC/E.coli polar lipid | PC/PA | PC/E.coli polar lipid | | PC/PA | |
| Supplemented ligands | ATP, $Mg^{2+}$, Hoechst | ATP, $Mg^{2+}$, Hoechst | ATP, $Mg^{2+}$ | ATP, $Mg^{2+}$ | | ATP, $Mg^{2+}$, Hoechst, Vanadate | |
| ***Conformational state*** | **BmrCD_IF-2H/ATP** | **BmrCD_IF-H/ATP** | **BmrCD_IF-ATP** | **BmrCD_IF-ATP2** | **BmrCD_OC-ATP** | **BmrCD_OC-ADPVi** | **BmrCD_IF-H/ADPVi** |
| PDB ID | 8FMV | 8SZC | 8FPF | 8T3K | 8FHK | 8T1P | ** |
| EMDB ID | EMD-29297 | EMD-40908 | EMD-29362 | EMD-41004 | EMD-29087 | EMD-40974 | EMD-41058 |
| Symmetry imposed | C1 | C1 | C1 | C1 | | C1 | |
| Microscope | Titan Krios (FEI) | Titan Krios (FEI) | Titan Krios (FEI) | Titan Krios (FEI) | | Titan Krios (FEI) | |
| Detector | Gatan K3 | Gatan K3 | Gatan K3 | Gatan K3 | | Gatan K3 | |
| Nominal magnification | 105,000 x | 105,000 x | 81,000 x | 130,000 x | | 130,000 x | |
| Voltage (kV) | 300 | 300 | 300 | 300 | | 300 | |
| Electron exposure (e/Å$^2$ ) | 48 | 51 | 54 | 52 | | 56 | |
| Defocus range (μm) | -0.8 to -2.2 | -0.8 to -2.2 | -0.8 to -1.6 | -0.4 to -2.2 | | -0.9 to -2.0 | |
| Pixel size (Å) | 0.818 | 0.818 | 1.1 | 0.647 | | 0.647 | |
| Number of Micrographs | 5696 | 12,706 | 3722 | 9114 | | 11,980 | |
| Particles images | 2,461,748 | 4,586,278 | 2,429,885 | 2,276,125 | 4,811,558 | 6,565,035 | 6,824,865 |
| Final particles images | 133,280 | 107,639 | 255,381 | 76,190 | 91,128 | 185,953 | 262,341 |
| Map resolution (Å) (FSC threshold=0.143) | 3.3 | 3.1 | 3.3 | 3.3 | 2.9 | 3.1 | 5.7 |
| **Refinement** | | | | | | | |
| Model resolution (Å) (original map, FSC threshold=0.5) | 3.6 | 3.7 | 3.8 | 7.1 | 3.8 | 3.2 | |
| Model resolution (Å) (composite map, FSC threshold=0.5) | 3.2 | 3.1 | 3.4 | 3.6 | 3.4 | -* | |
| B-factor used for map sharpening (Å$^2$ ) | -104.6 | -77.9 | -112.4 | -86.1 | -90.7 | -89.5 | |
| ***Model composition*** | | | | | | | |
| Non-hydrogen atoms | 10,501 | 10,511 | 10,624 | 10,144 | 10,120 | 10,587 | |
| Protein residues | 1236 | 1235 | 1,226 | 1,220 | 1,221 | 1,241 | |
| ATP, Hoechst, lipids, $Mg^{2+}$, AOV | 2, 2, 20, 1, 0 | 2, 1, 21, 1, 0 | 2, 0, 25, 0, 0 | 2, 0, 13, 0, 0 | 2, 0, 16, 2, 0 | 1, 0, 21, 2, 1 | |
| ***Mean B factors (Å$^2$)*** | | | | | | | |
| Protein | 78.26 | 70.63 | 79.69 | 62.42 | 61.64 | 59.73 | |
| Ligands | 75.14 | 72.92 | 63.71 | 50.26 | 60.04 | 77.21 | |
| ***R.m.s. deviations*** | | | | | | | |
| Bond lengths (Å) | 0.014 | 0.004 | 0.008 | 0.005 | 0.008 | 0.006 | |
| Bond angles (°) | 1.422 | 0.695 | 0.895 | 0.769 | 1.136 | 0.782 | |
| ***Molprobity score*** | 1.73 | 1.74 | 1.77 | 1.89 | 1.86 | 1.96 | |
| ***Clash score*** | 5.92 | 5.48 | 5.93 | 8.02 | 7.55 | 8.22 | |
| ***Poor rotamers (%)*** | 0.10 | 0.00 | 0.00 | 0.10 | 0.39 | 0.38 | |
| ***Ramachandran plot*** | | | | | | | |
| Favored (%) | 93.89 | 93.00 | 93.10 | 92.84 | 93.04 | 91.34 | |
| Allowed (%) | 6.11 | 7.00 | 6.90 | 7.16 | 6.96 | 8.66 | |
| Disallowed (%) | 0.00 | 0.00 | 0.00 | 0.00 | 0.00 | 0.00 | |

*: The model of BmrCD_OC-ADPVi was built by original map after Local refinement in CryoSPARC.
**: No structural model was built for BmrCD_IF-H/ADPVi.

### Supplementary Table 2: Root mean square deviation (r.m.s.d.) of overall structure.

| | BmrCD_det | BmrCD_IF-2H/ATP | BmrCD_IF-H/ATP | BmrCD_IF-ATP | BmrCD_IF-ATP2 | BmrCD_OC-ATP | BmrCD_OC-ADPVi |
|---|---|---|---|---|---|---|---|
| **BmrCD_det** | 0.00 | 0.65 (959 Cα) | 0.82 (992 Cα) | 1.38 (1078 Cα) | 0.92 (980 Cα) | 6.24 (1190 Cα) | 6.30 (1217 Cα) |
| **BmrCD_IF-2H/ATP** | | 0.00 | 0.42 (1131 Cα) | 0.94 (1153 Cα) | 0.67 (1026 Cα) | 6.48 (1199 Cα) | 6.48 (1228 Cα) |
| **BmrCD_IF-H/ATP** | | | 0.00 | 0.87 (1142 Cα) | 0.75 (1031 Cα) | 6.60 (1202 Cα) | 6.58 (1229 Cα) |
| **BmrCD_IF-ATP** | | | | 0.00 | 0.99 (1096 Cα) | 6.64 (1193 Cα) | 6.62 (1219 Cα) |
| **BmrCD_IF-ATP2** | | | | | 0.00 | 6.46 (1172 Cα) | 6.40 (1194 Cα) |
| **BmrCD_OC-ATP** | | | | | | 0.00 | 0.63 (1039 Cα) |
| **BmrCD_OC-ADPVi** | | | | | | | 0.00 |

Superposition was performed at the residue level by Pymol and r.m.s.d. was calculated at the Cα level in unit of Å.

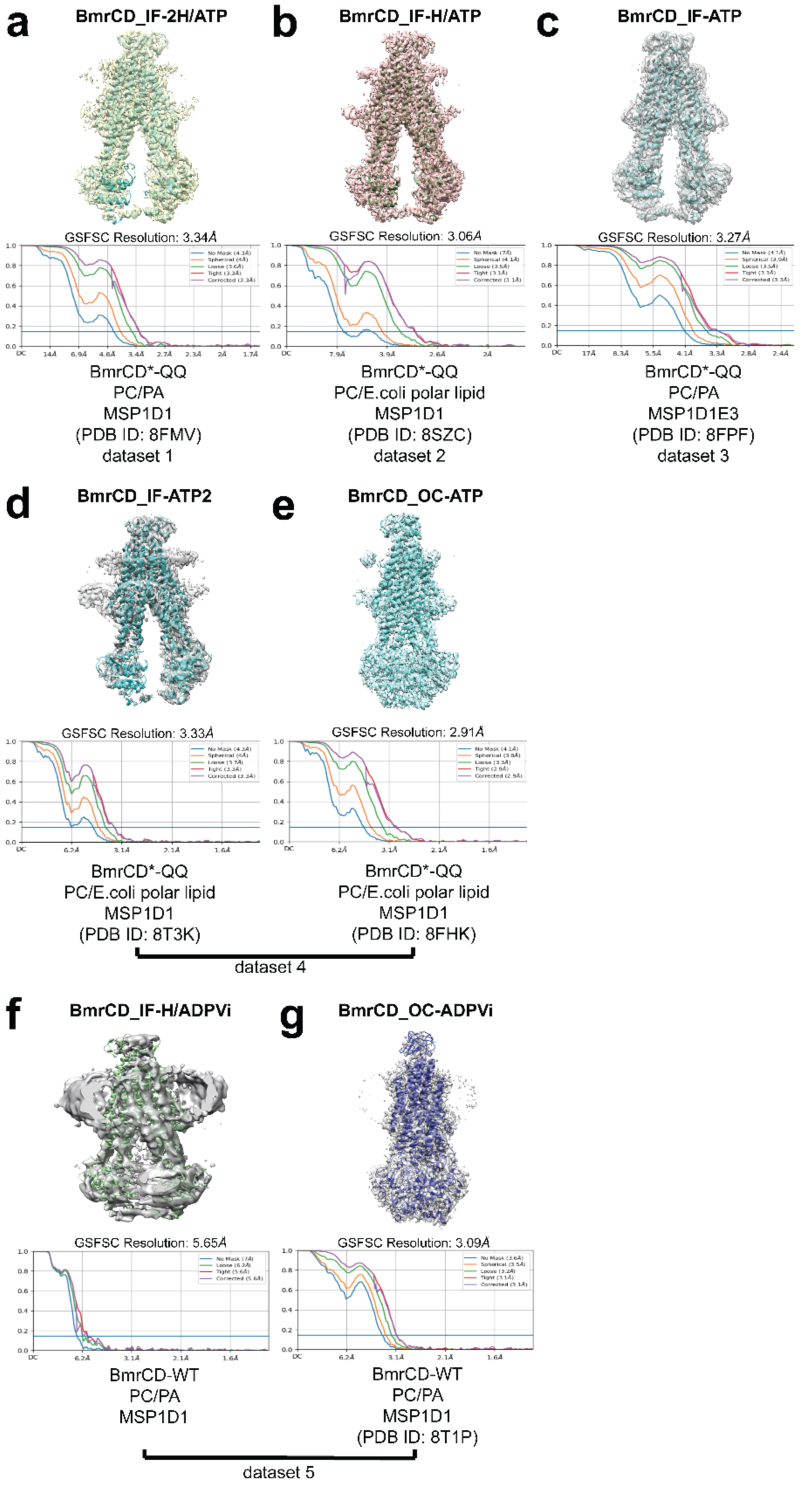


**Supplementary Fig. 1: Conformational gallery of BmrCD.**
Seven cryo-EM density maps from 5 datasets are shown, and the cartoon models are fitted into the maps (For more information see Methods). Original maps estimated resolution is based on Fourier shell correlation (FSC) curves, **a**-**e** are original maps after

local refinement in cryoSPARC; **f** and **g** are original maps after NU refinement in cryoSPARC. The BmrCD constructs, lipid composition of the nanodiscs and MSP used for reconstitution are shown below the maps. BmrCD*-QQ and BmrCD-WT refer to cysteine-less BmrCD in QQ background and wild-type BmrCD respectively.

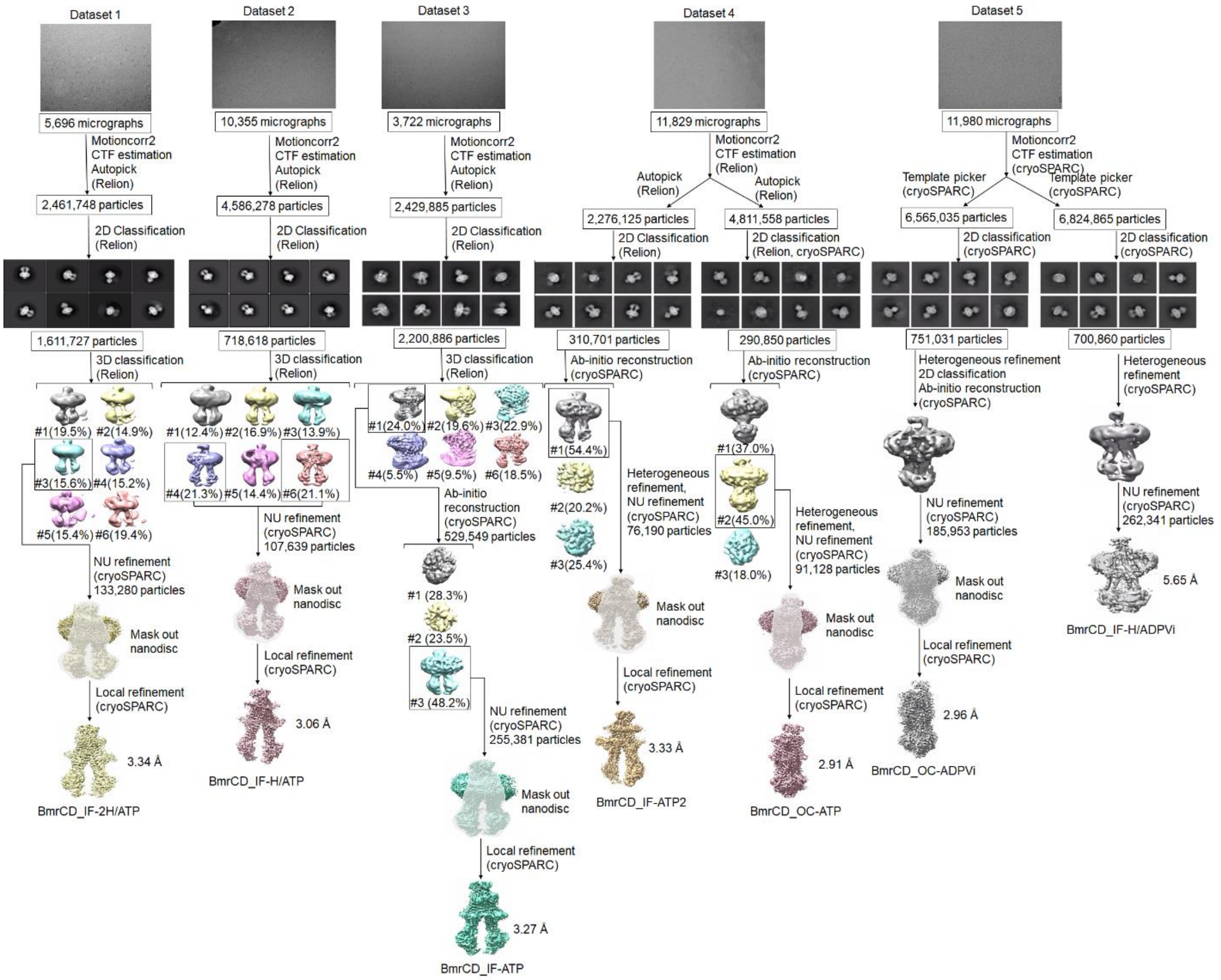


**Supplementary Fig. 2: Cryo-EM data processing workflow for BmrCD datasets.** The diagram schematically depicts all the five BmrCD cryo-EM datasets that were collected in this study and the computational processing steps used to reconstruct the seven BmrCD original maps: BmrCD_IF-2H/ATP, BmrCD_IF-H/ATP, BmrCD_IF-ATP, BmrCD_IF-ATP2, BmrCD_OC-ATP, BmrCD_OC-ADPVi, and BmrCD_IF-H/ADPVi. Programs used for each step are noted, and the representative micrographs, 2D and 3D images are shown above.

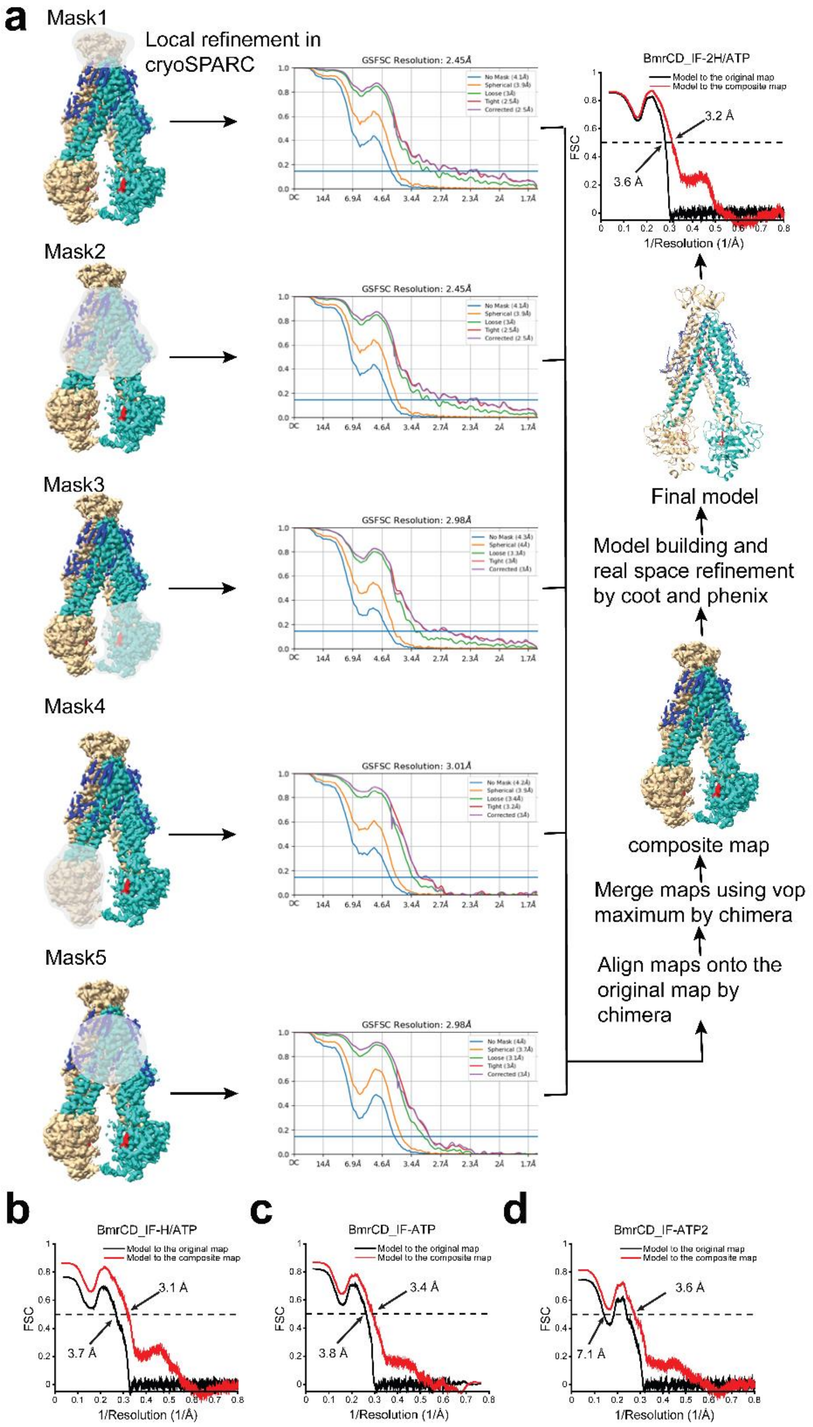


**Supplementary Fig. 3: Diagram showing the local refinement strategy for BmrCD inward facing conformations.**

**a** Diagram showing the local refinement strategy for BmrCD_IF-2H/ATP. Masks are shown transparent, and subunits colored as in Figure 1. Fourier shell correlation (FSC) curves for each mask after local refinement in CryoSPARC are shown. FSC curves of the refined model versus the original map after NU refinement in cryoSPARC (black) and the composite map (red) are shown at the final step in the diagram. **b**, **c**, and **d** Similar local refinement strategy was applied and the FSC curves of the refined model versus the original map (black) and the composite map (red) are plotted for BmrCD_IF-H/ATP (**b**), BmrCD_IF-ATP (**c**), BmrCD_IF-ATP2 (**d**).

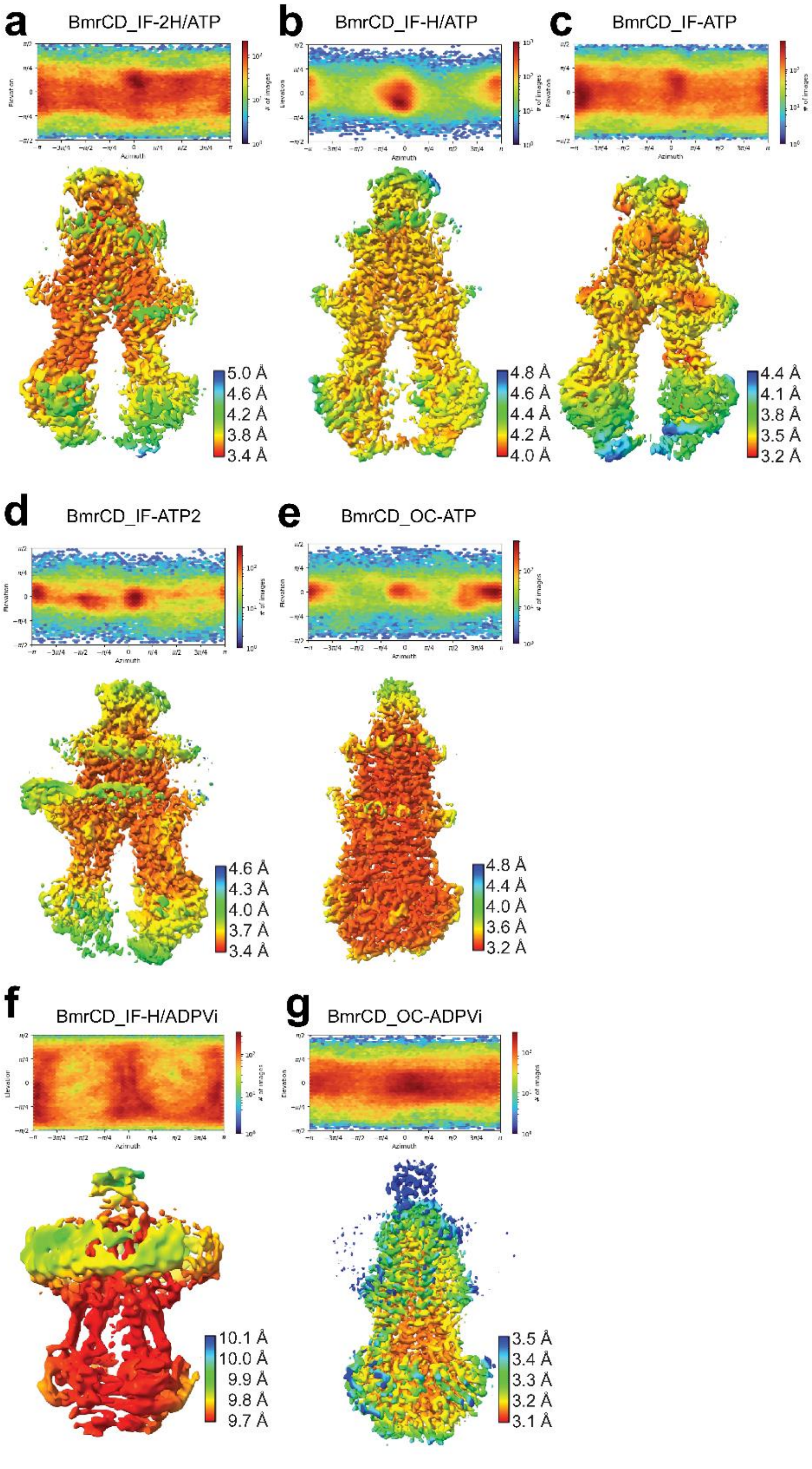


**Supplementary Fig. 4: Angular distribution of particles and local resolution maps for BmrCD.**

**a**-**e**, and **g** angular distribution of particles (upper panel) and local resolution map (lower panel) for original maps after local refinement in cryoSPARC for BmrCD_IF-2H/ATP (**a**),

BmrCD_IF-H/ATP (**b**), BmrCD_IF-ATP (**c**), BmrCD_IF-ATP2 (**d**), BmrCD_OC-ATP (**e**), BmrCD_OC-ADPVi (**g**). **f** angular distribution of particles (upper panel) and local resolution map (lower panel) for original maps after NU refinement in cryoSPARC for BmrCD_IF-H/ADPVi.

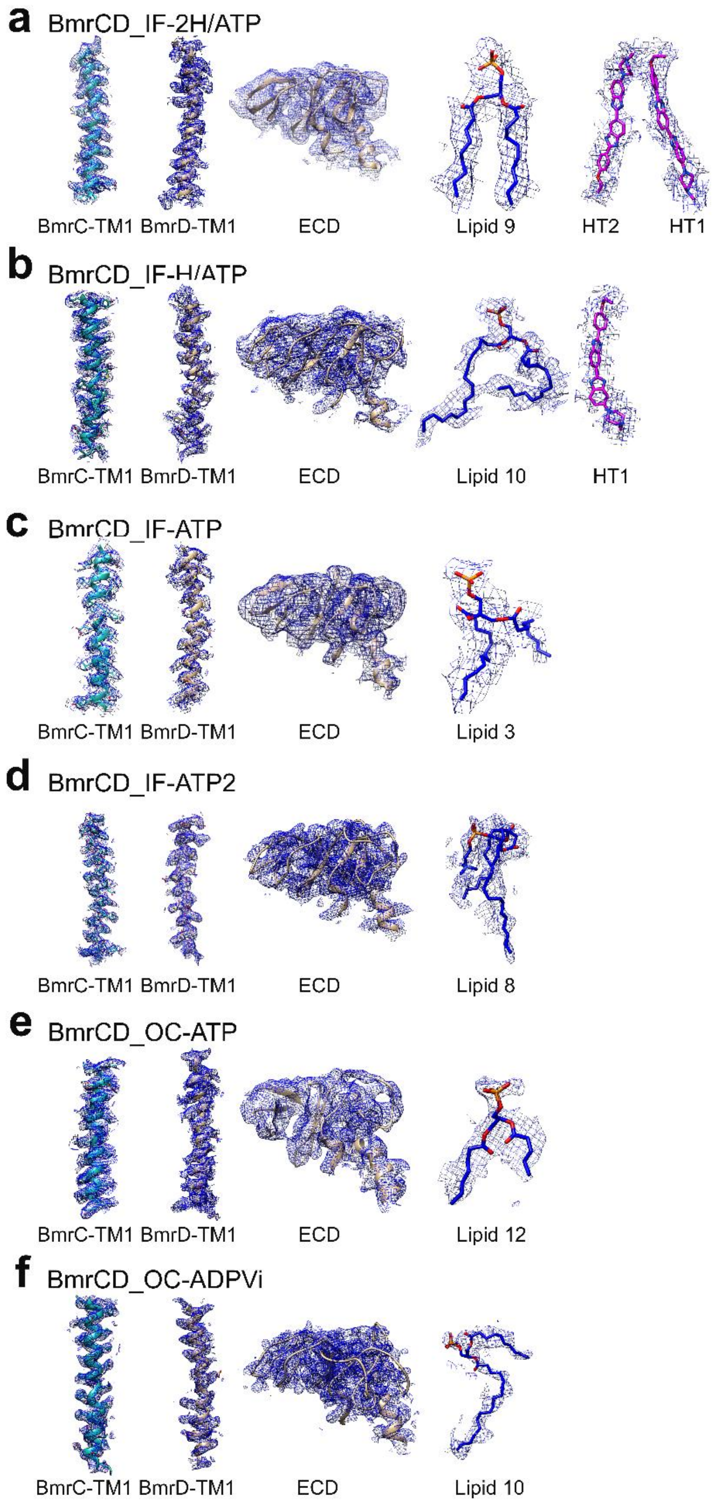


**Supplementary Fig. 5: Representative density maps for BmrCD.**

Transmembrane domain helices TM1 of BmrC and BmrD are selected for illustration of the clear side-chain densities. Extracellular domains (ECD), selected lipids molecules (labeled with their ID in the structures) and substrates Hoechsts (HT1 and HT2) are shown along with their respective densities.

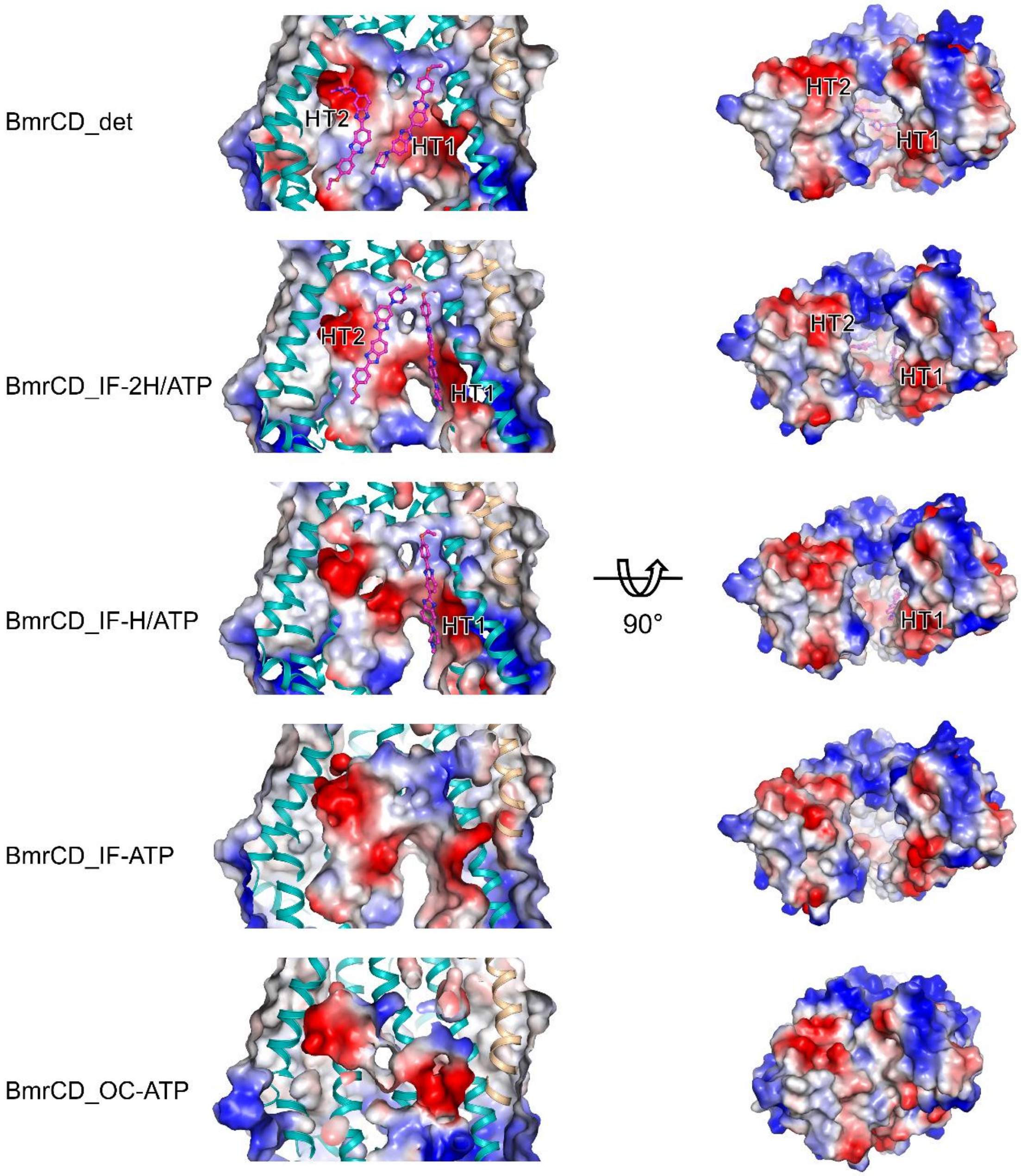


**Supplementary Fig. 6: Electrostatics analysis of Hoechst-binding pocket.**

Upper panels show side views of the electrostatic surface of the substrate Hoechst-binding pocket of BmrCD_det, BmrCD_IF-2H/ATP, BmrCD_IF-H/ATP, BmrCD_IF-ATP, and BmrCD_OC-ATP. The lower panels show the bottom view of the TMD. The surfaces are colored from blue (positively charged regions) to red (negatively charged regions). Neutral regions are shown as white. BmrCD is shown in cartoon and Hoechst is shown in stick. Two Hoechst molecules are labeled as HT1 and HT2.

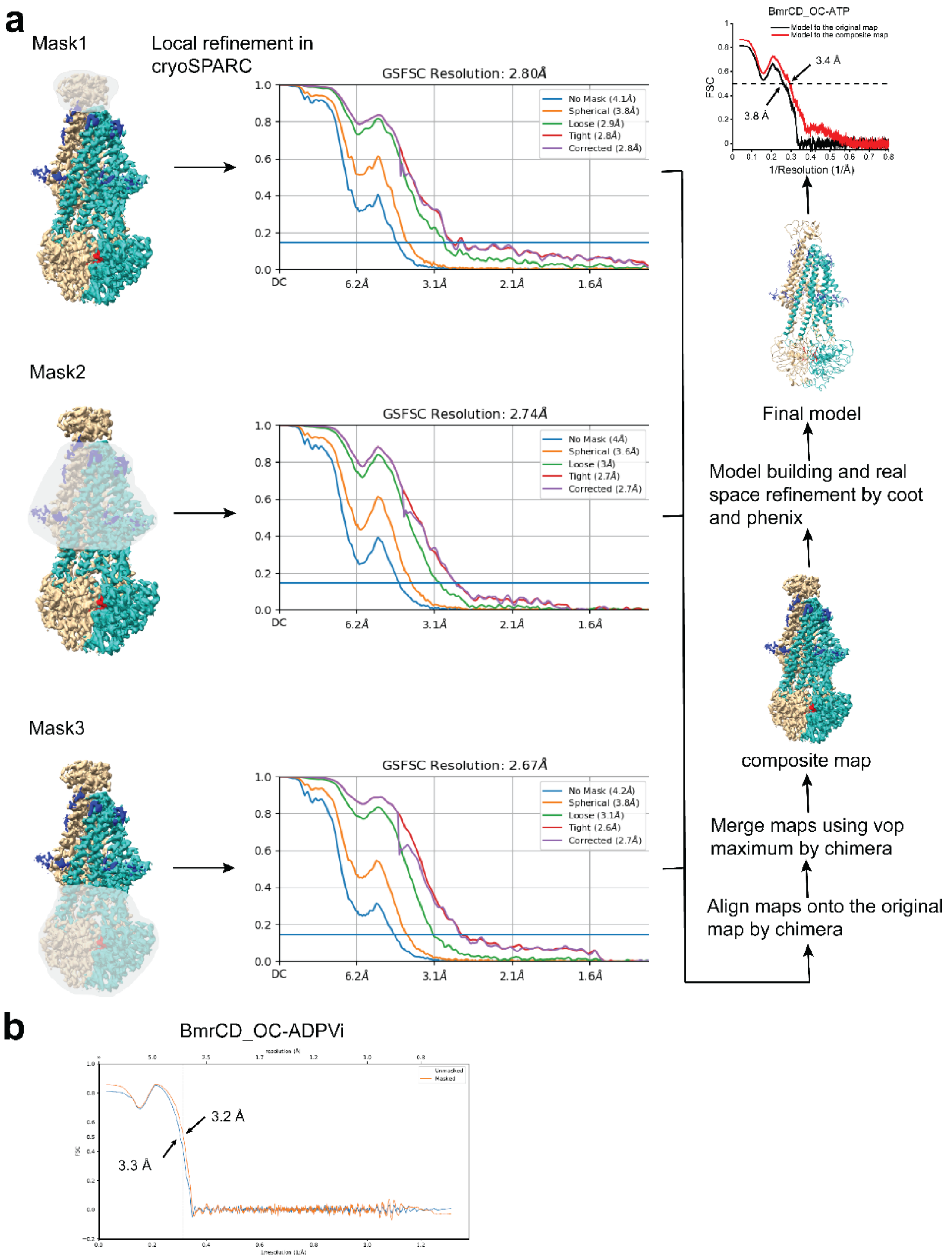


**Supplementary Fig. 7: Refinement strategies for BmrCD_OC-ATP and BmrCD_OC-ADPVi.**

**a** Local refinement strategy for BmrCD_OC-ATP. Masks are shown transparent, and subunits colored as in Figure 1. Fourier shell correlation (FSC) curves for each mask after local refinement in CryoSPARC are shown. FSC curves of the refined model versus the original map after NU refinement in cryoSPARC (black) and the composite map (red) are shown at the final step in the diagram. **b** Real-space refinement for BmrCD_OC-ADPVi. Fourier shell correlation curves show model fit after Real-space refinement for BmrCD_OC-ADPVi. The local refinement strategy to obtain a composite map was not applied for BmrCD_OC-ADPVi (See Supplementary Table 1 and Figure S2).

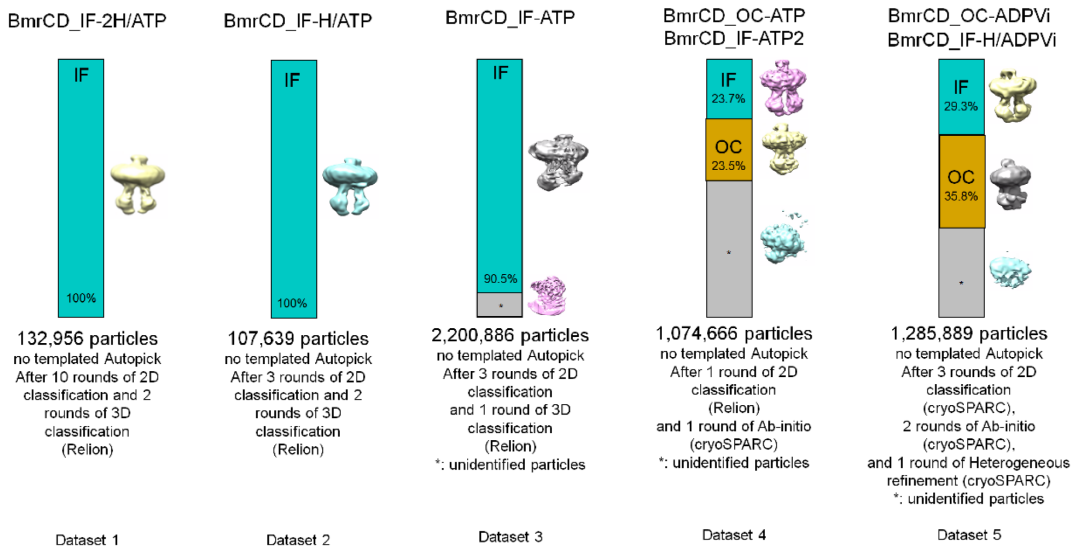


**Supplementary Fig. 8: Particle distributions in the 5 datasets.**
The percentage of the particles' distributions are listed, and the representative image of the conformations are shown.

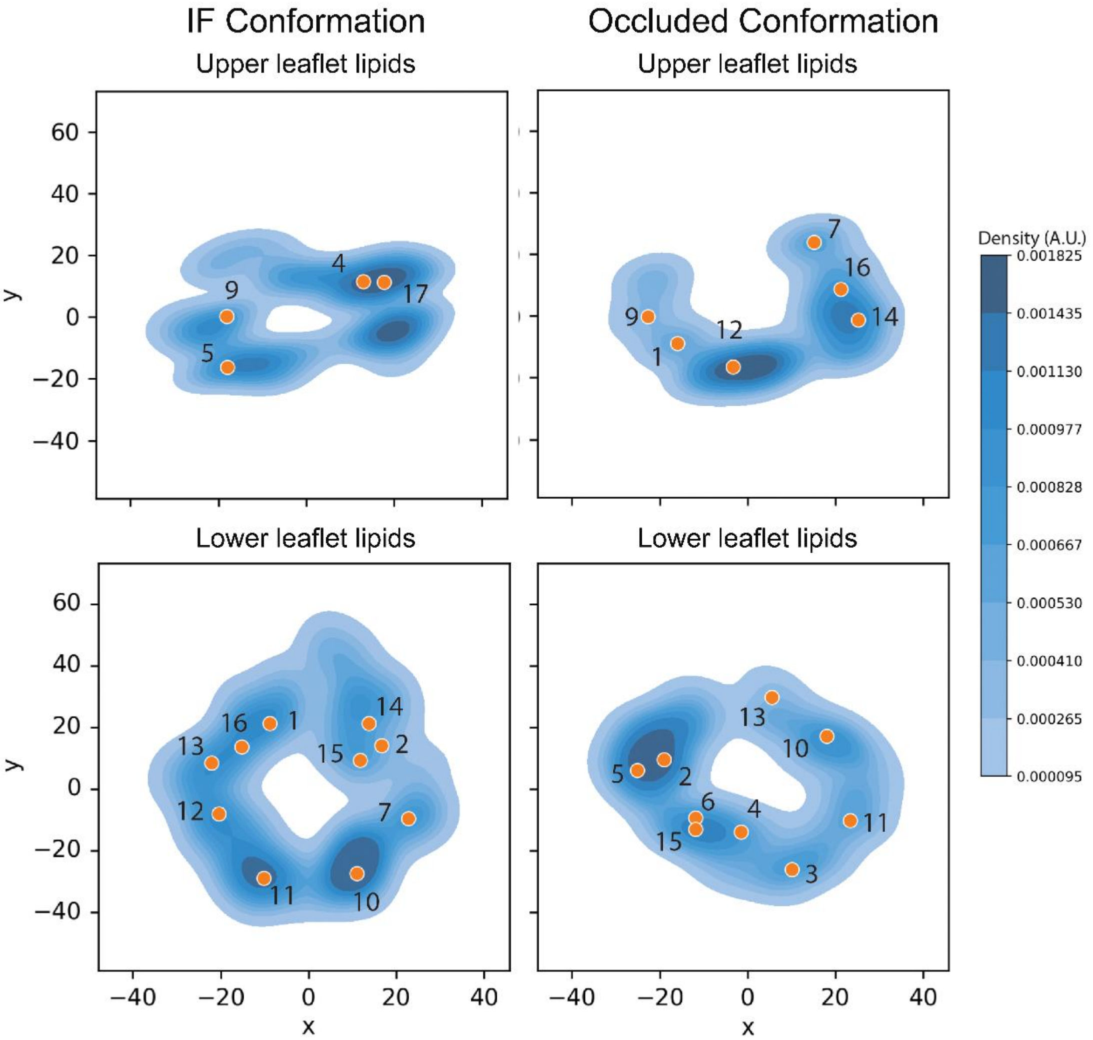


**Supplementary Fig. 9: Lipids distributions in BmrCD cryo-EM structures.** Kernel density estimation was used to plot the lipids distribution in BmrCD cryo-EM structures excluding lipids without observable headgroup. The lipid positions of phosphorus atoms from BmrCD_IF-2H/ATP are marked in orange dots with numbers for reference while those for BmrCD_IF-H/ATP, BmrCD_IF-ATP, and BmrCD_IF-ATP) are represented by the lipid density. Similar lipid density for the two occluded conformations (BmrCD_OC-ATP and BmrCD_OC-ADPVi) are plotted. The lipid positions of phosphorus atoms from BmrCD_OC-ATP are marked in orange dots and numbers for reference. The regions with oval darker blue densities highlight conserved lipid binding sites (lipids 4, 17, 10, and 11 in IF conformation; 12, 14, 2, and 5 in occluded conformation), and the regions with smeared densities nearby indicate less conserved lipid binding sites in the cryo-EM structures.

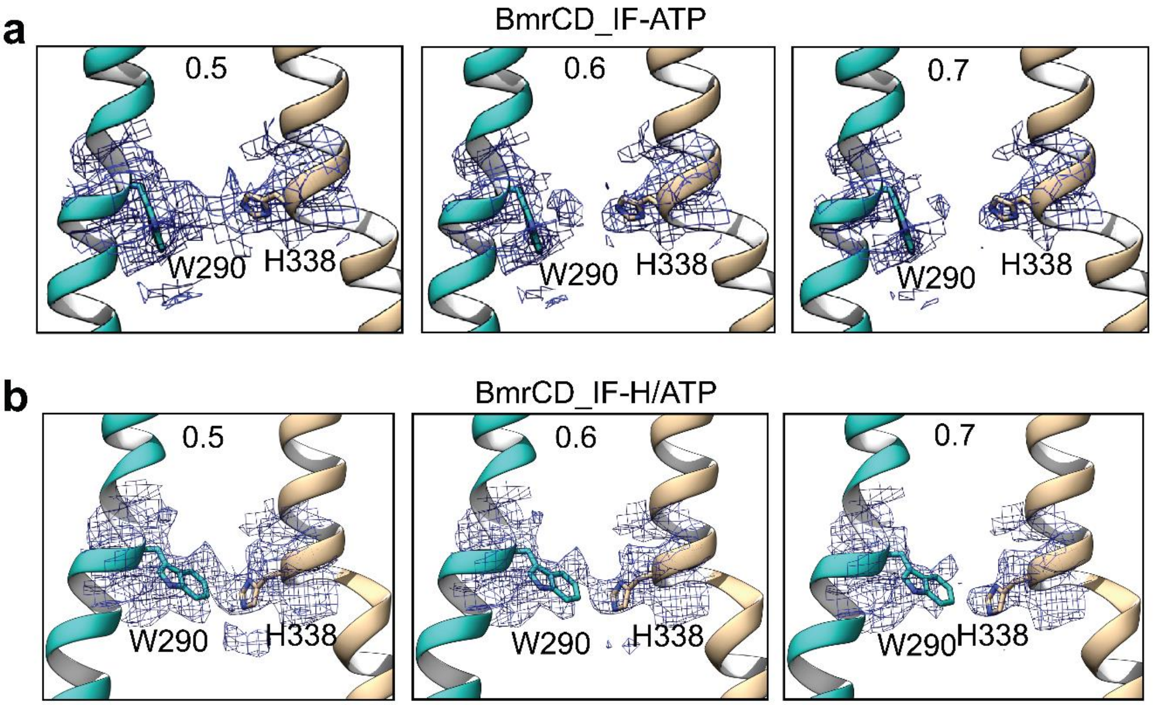


**Supplementary Fig. 10: Density maps of WH-latch in BmrCD_IF-ATP and BmrCD_IF-H/ATP.**
The density maps are contoured at three contour levels as indicated. The density of W290 in BmrCD_IF-ATP (**a**) indicates that W290 may also adopt a locked latch conformation in BmrCD_IF-H/ATP (**b**) besides the major open conformation to block HT binding.

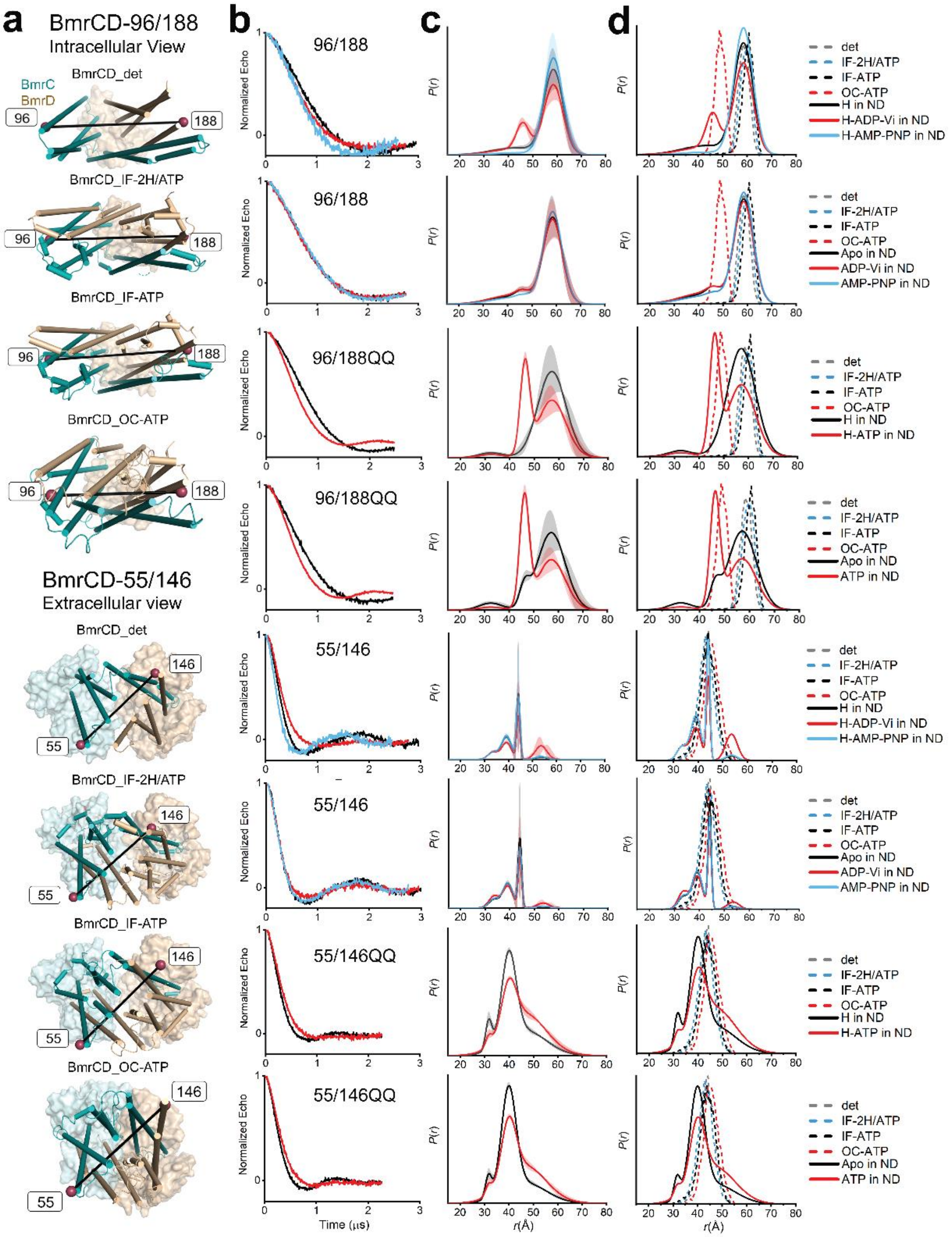


**Supplementary Fig. 11: DEER decay signals for spin-labeled BmrCD mutants in the TMD.**

**a** Cartoon representations of BmrCD highlighting the spin-labeled positions. **b** Normalized Echo decay intensity curves. **c** Distance distribution analyzed as described in the methods. The light color bands represent confidence bands. **d** Distance distribution from DEER (solid lines) compared with predicted distribution from cryo-EM structures (dashed lines).

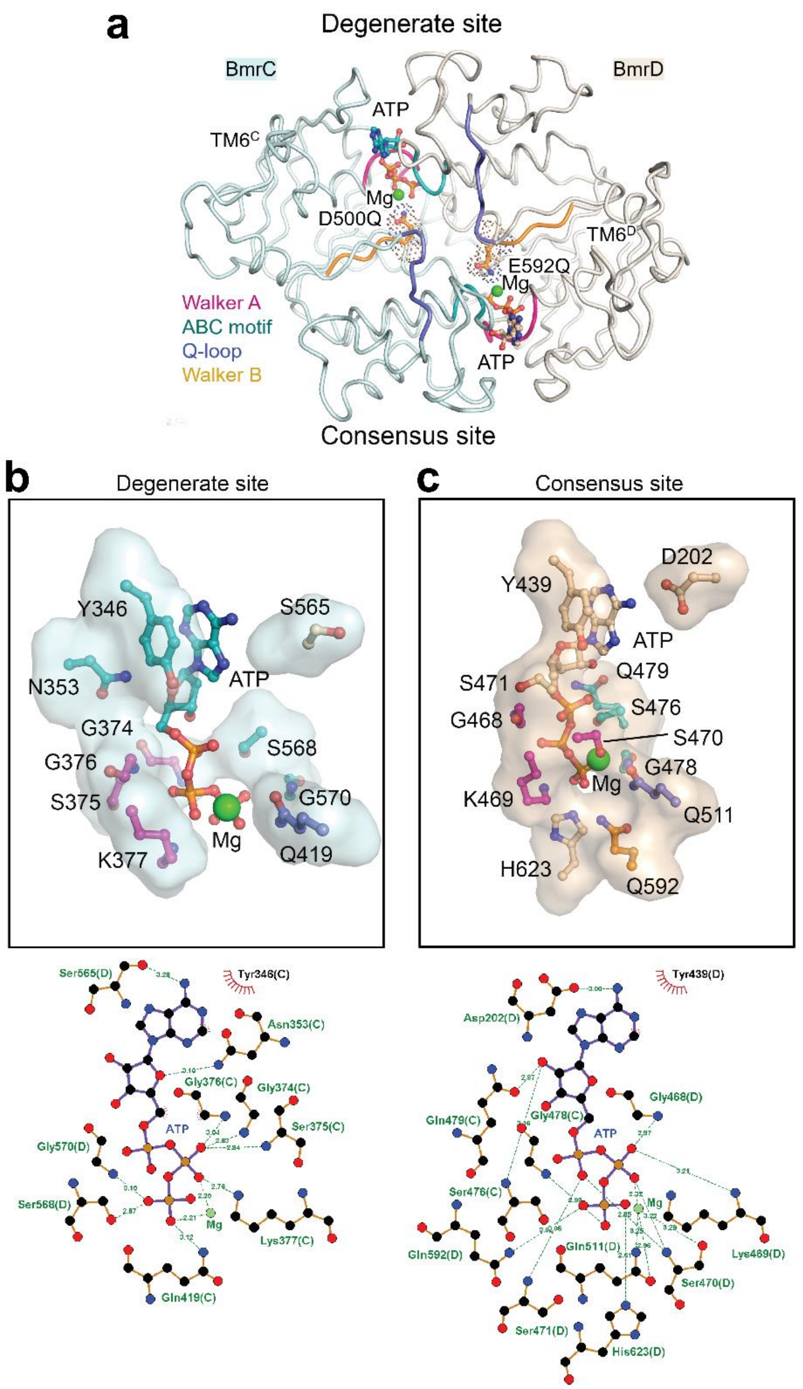


**Supplementary Fig. 12: Asymmetric coordination of ATP in the OC conformation (BmrCD_OC-ATP).**

**a** Close-up views of the degenerate and consensus NBSs. BmrCD is shown in ribbon representation with catalytic residues side chains and ATP represented by sticks. Conserved motifs are highlighted. **b** and **c** The atomic features of the complete ATP-binding pockets in the degenerate NBS (**b**), and the consensus NBS (**c**). The key residues from the conserved motifs are shown in sticks in the same colors, and LigPlot schematics of ATP binding are shown below. Interactions between ATP and BmrCD_OC-ATP at the degenerate and consensus NBSs were plotted by LigPlot(Laskowski and Swindells, 2011) with default hydrogen-bond calculation parameters setting (Maximum H-A distance is set as 2.70 Å and maximum D-A distance is set as 3.35 Å). Electrostatic and hydrophobic interactions are shown in dashes and eyelashes, respectively.

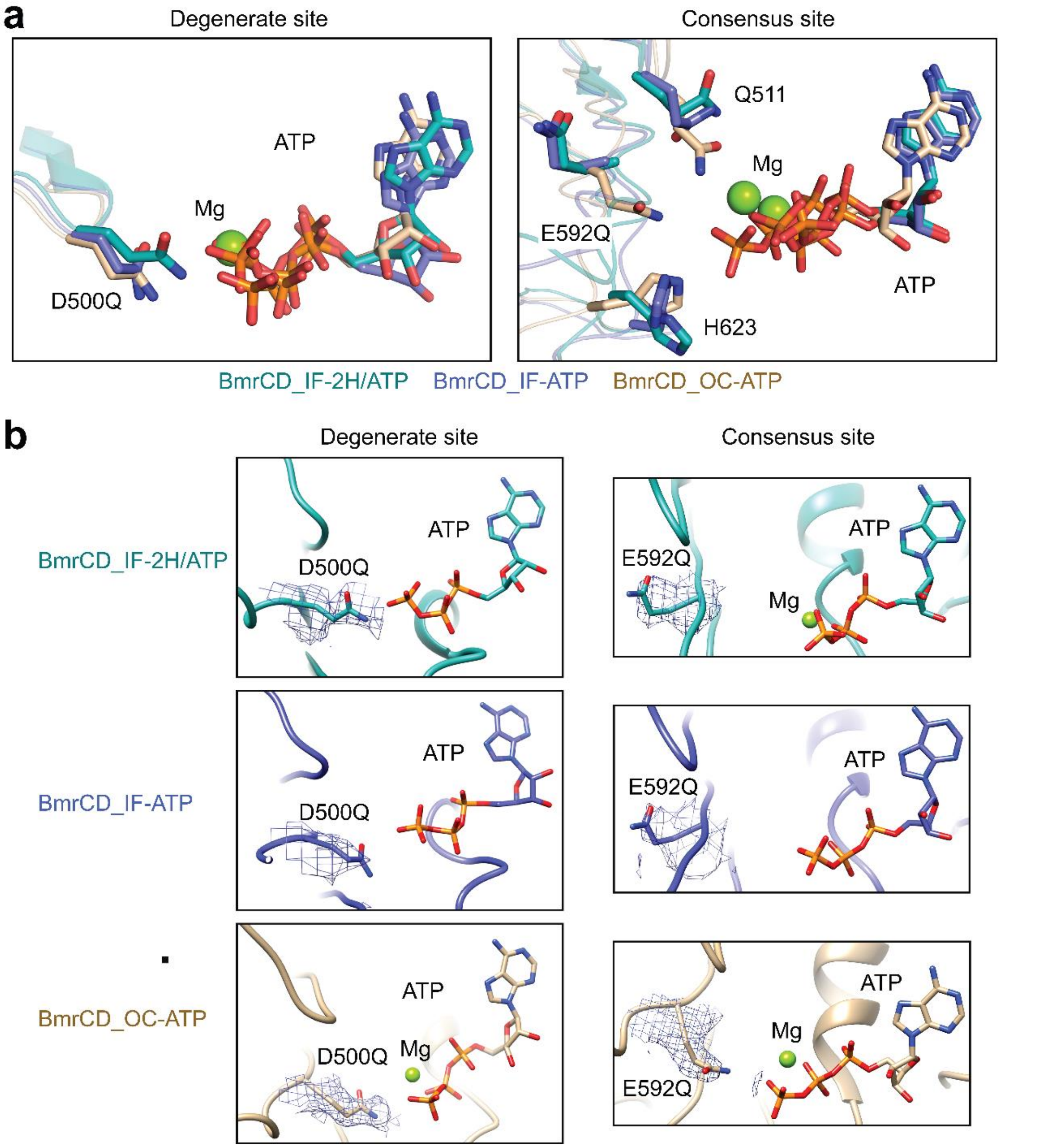


**Supplementary Fig. 13: Configurations of catalytic residues in the NBSs.**

**a** Superimposition of BmrCD NBDs to highlight the catalytic residues D500Q and E592Q in degenerate NBS and consensus NBS, respectively. **b** Density maps of the catalytic residues.

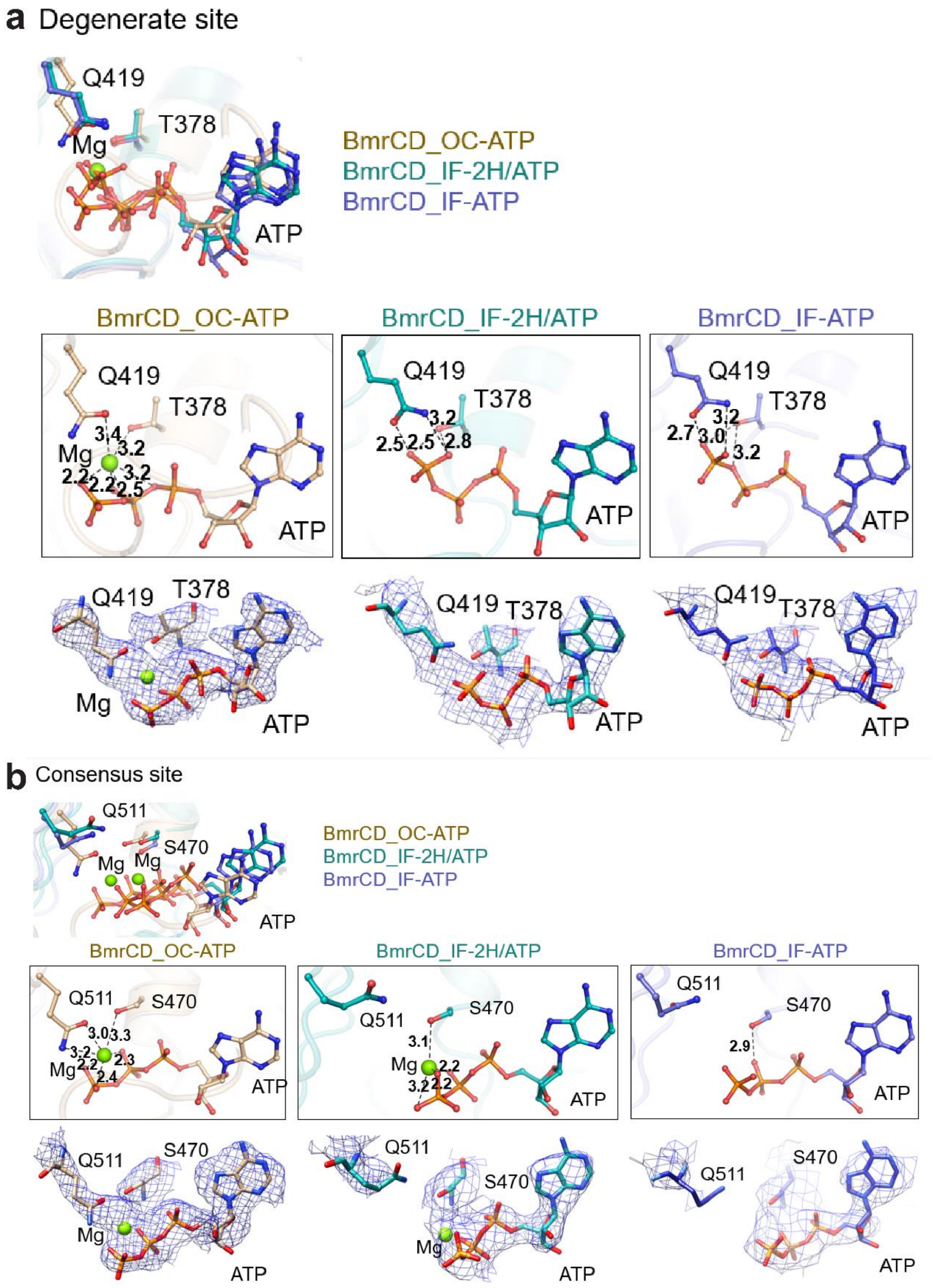


**Supplementary Fig. 14: Magnesium ions in BmrCD conformations.**

**a** Coordination of $Mg^{2+}$ in the degenerate NBS. The distance of the $Mg^{2+}$ with its coordinated residues and relative distances (Å) are shown in boxed pictures and density maps are shown below. Superimposition of the coordination sites are shown above.
**b** Similar analysis of the coordination of $Mg^{2+}$ in the consensus NBS.

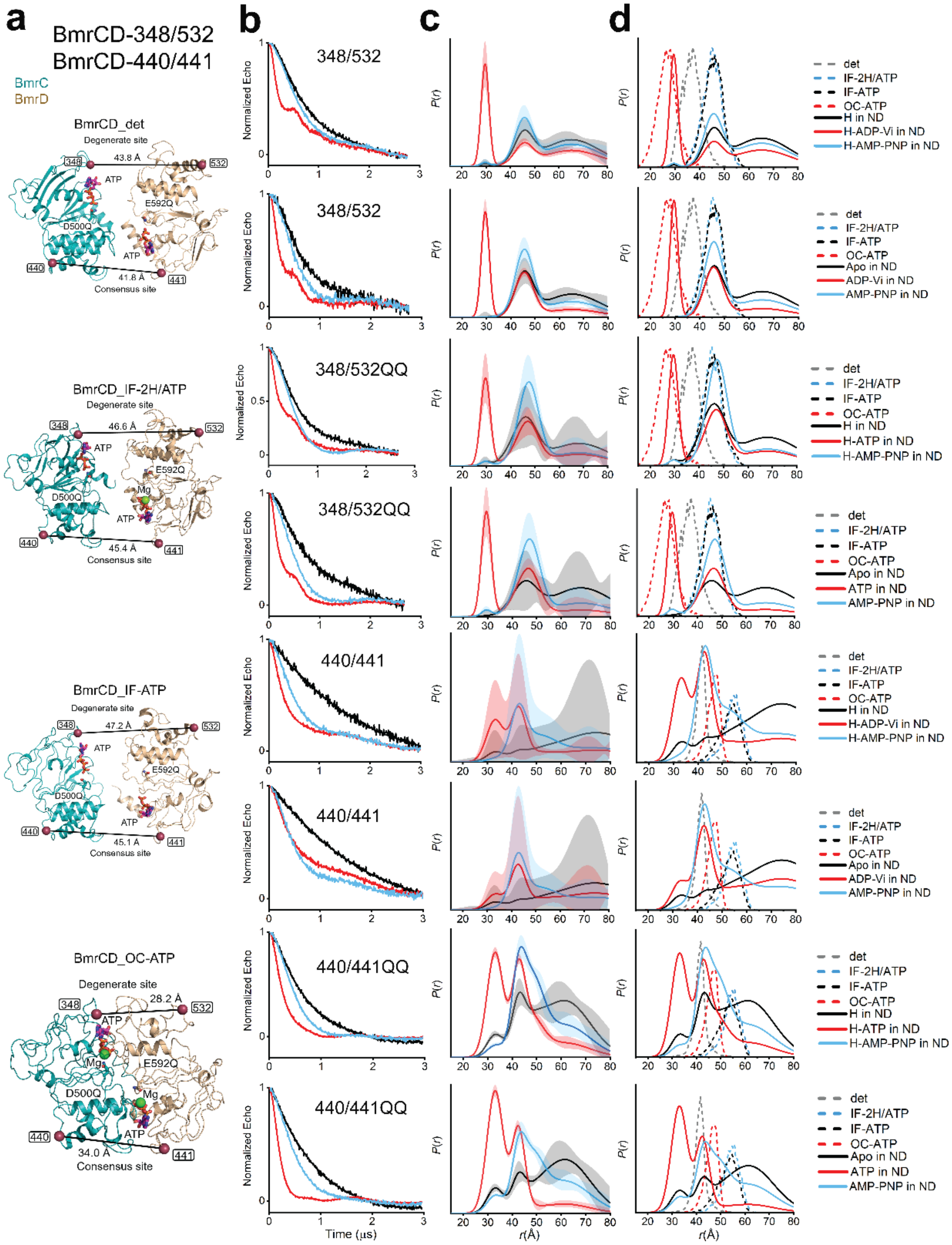


**Supplementary Fig. 15: DEER decay signals for spin-labeled BmrCD mutants in the NBD.**

**a** Cartoon representations of BmrCD highlighting the spin-labeled positions. **b** Normalized Echo decay intensity curves. **c** Distance distribution analyzed as described in the methods. The light color bands represent confidence bands. **d** Distance distribution from DEER (solid lines) compared with predicted distribution from cryo- EM structures (dashed lines).

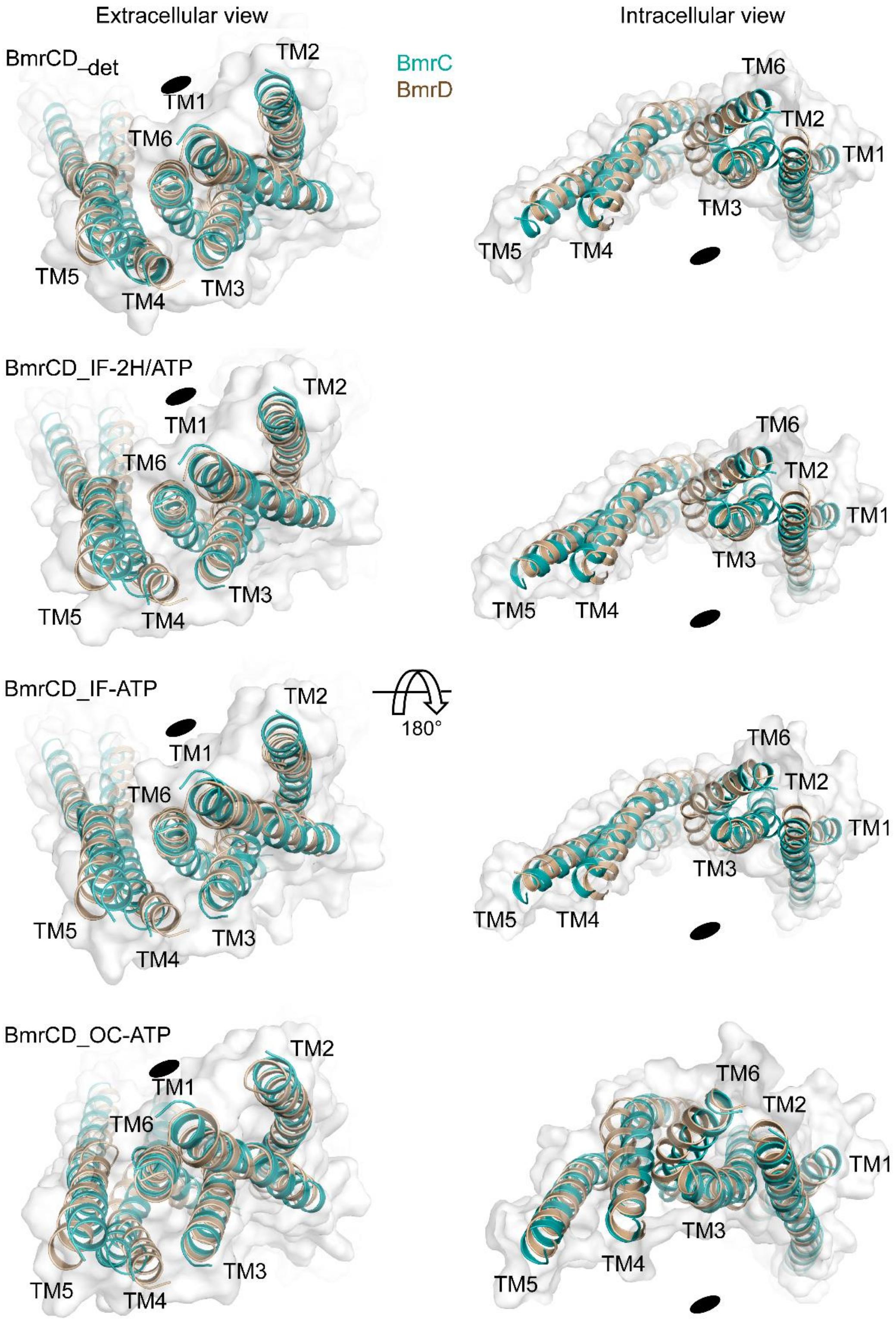


326
327 **Supplementary Fig. 16: Superimposition of transmembrane helices from BmrC**
328 **and BmrD highlighting asymmetry.**

Transmembrane helices are shown in cartoon and labeled. The black ellipse represents the symmetry axis.

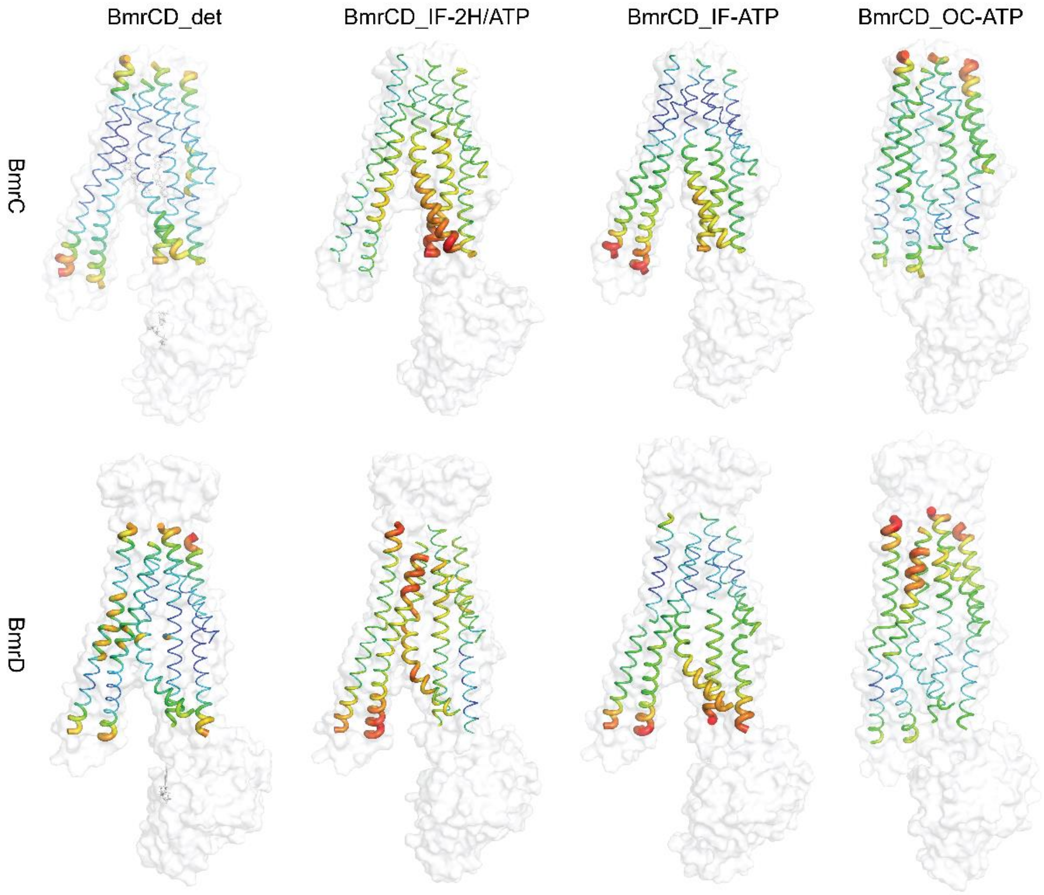


**Supplementary Fig. 17: B-factor maps of transmembrane helices.**

B-factor maps of the transmembrane helices of BmrC and BmrD are generated by “b factor putty” in PyMol.

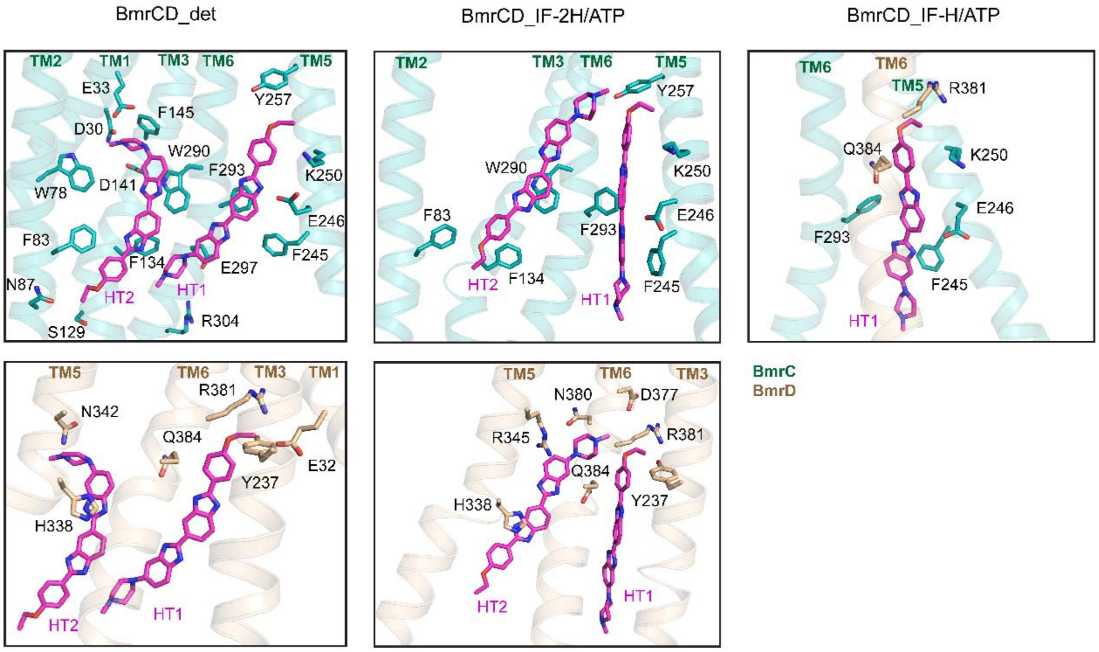


**Supplementary Fig. 18: Hoechst-binding pockets in BmrCD_det, BmrCD_IF-2H/ATP, and BmrCD_IF-H/ATP.**

Representative residues of the binding pocket of Hoechst (H) are shown as green/tan sticks. Substrate Hoechsts (HTs) are shown as magenta sticks. Upper panels and lower panels in BmrCD_det and BmrCD_IF-2H/ATP are in zoomed side view of the interactions with BmrC and BmrD, respectively.

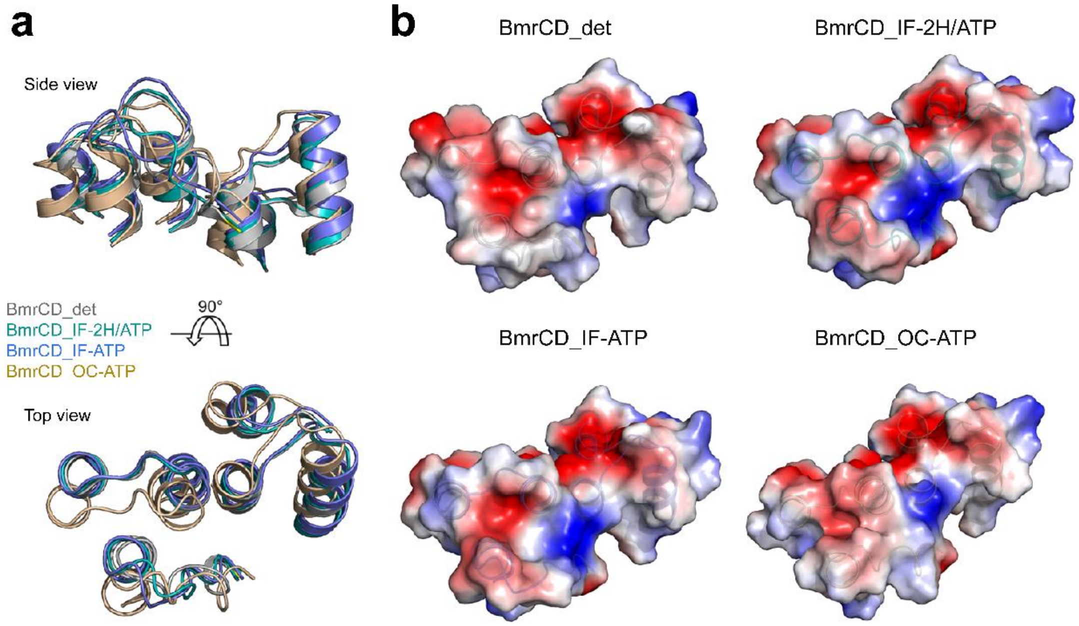


**Supplementary Fig. 19: The extracellular gate of BmrCD.**

**a** Superimposition of the extracellular gate in cartoon. **b** Electrostatics surface analysis of the extracellular gate of BmrCD.

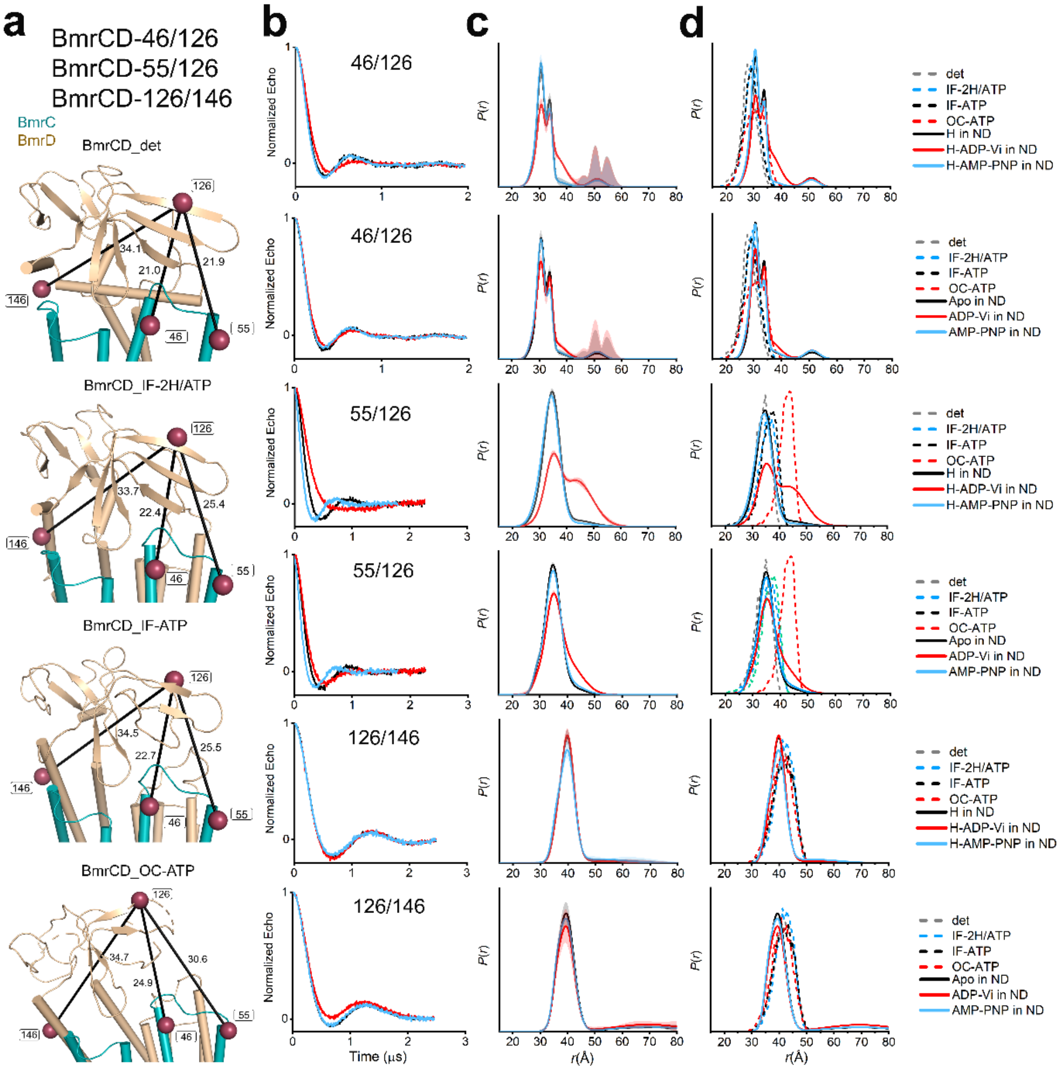


**Supplementary Fig. 20: DEER decay signals for spin-labeled BmrCD mutants in the ECD.**

**a** Cartoon representations of BmrCD highlighting the spin-labeled positions. **b** Normalized Echo decay intensity curves. **c** Distance distribution analyzed as described in the methods. The light color bands represent confidence bands. **d** Distance distribution from DEER (solid lines) compared with predicted distribution from cryo-EM structures (dashed lines).

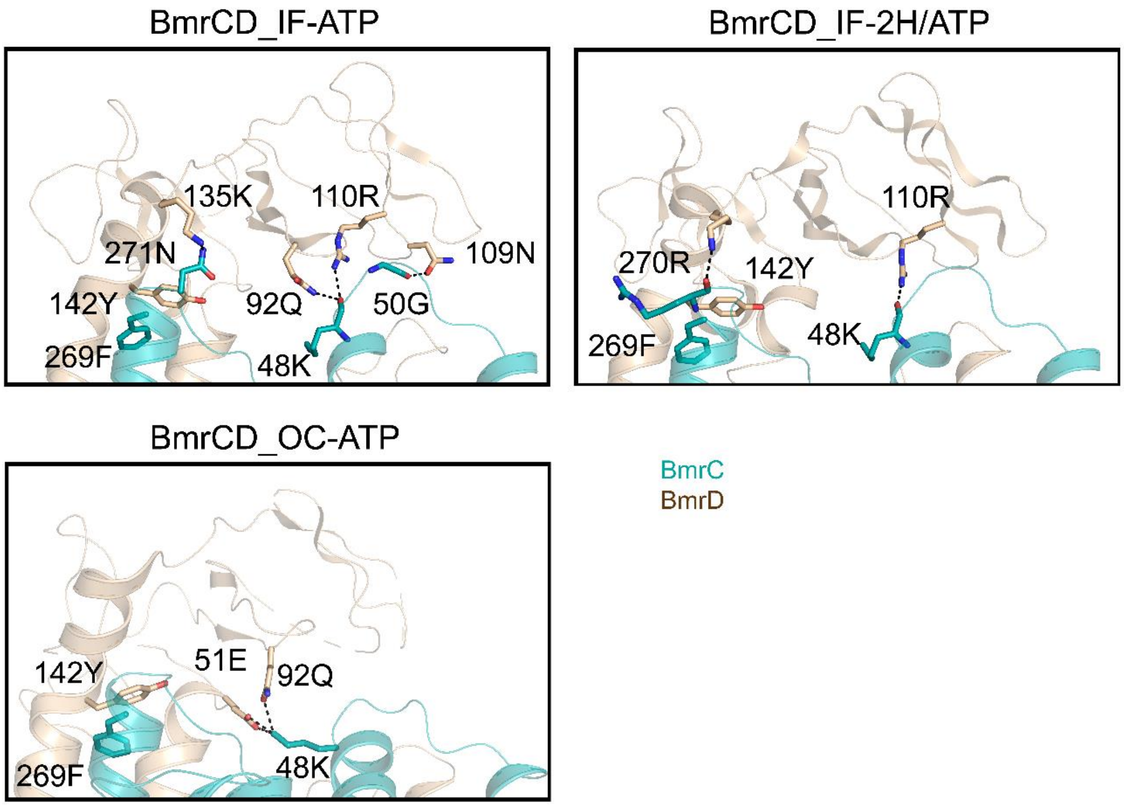


**Supplementary Fig. 21: Interactions between the ECD and TMD.**

Interactions between ECD and TMD of BmrCD_IF-2H/ATP, BmrCD_IF-ATP BmrCD_OC-ATP. The sidechains of the relative residues are represented by stick. Hydrogen bonds are shown in dashed line.

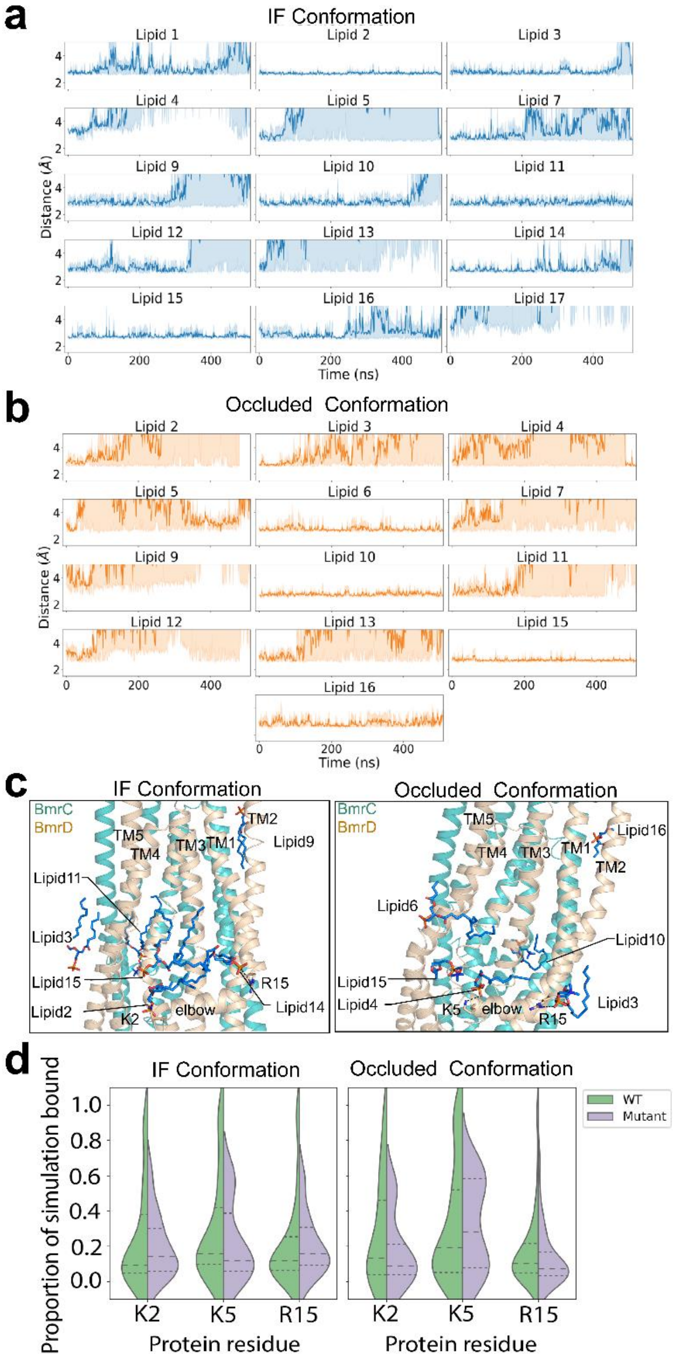


**Supplementary Fig. 22: Assessment of lipid binding by MD simulations.**
**a** and **b** Molecular dynamics (MD) simulation results of modeled lipids on BmrCD_IF-2H/ATP (**a**) and BmrCD_OC-ATP (**b**), respectively. Minimum distance between modeled lipids and protein over time averaged across all simulation replicas with error bars shown in light blue. These plots highlight stably bound lipids which preserve their position with respect to the protein across all simulations for each protein conformation. **c** Close-up view of TMDs shows stably binding lipids confirmed by the MD simulations. Lipids are shown in sticks with labeling. Possible interaction residues of BmrCD are represented in sticks, and hydrogen bonds are shown as dashed lines. **d** Lipid binding

residence time is shown as densities for both WT and mutant BmrCD systems. Comparison of these densities demonstrates a decrease in high-residence lipid binding for any of the BmrD elbow helix mutants K2A, K5A and R15A whose positions are shown in (**c**).

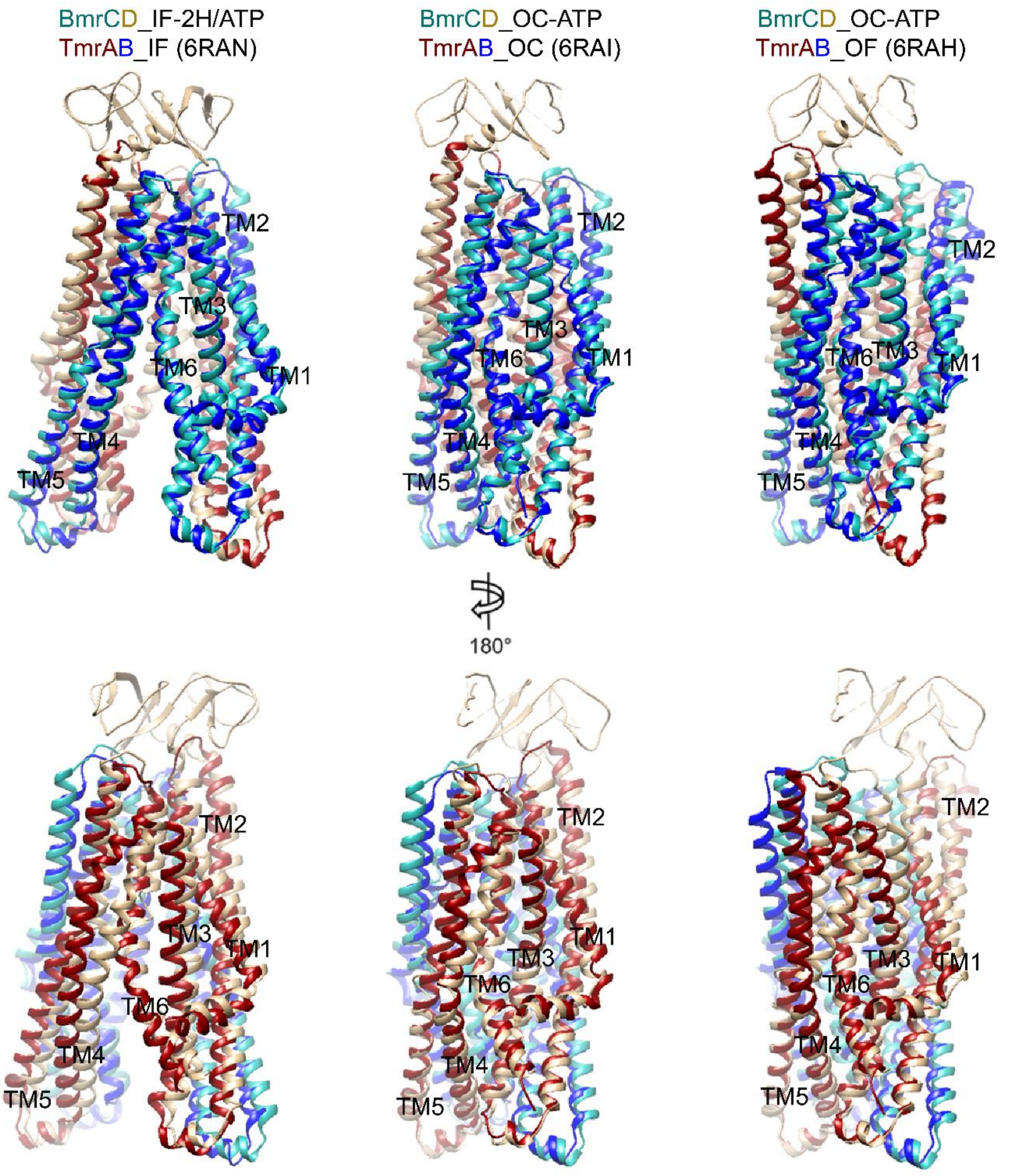


**Supplementary Fig. 23: BmrCD compared to TmrAB in different conformations.**
BmrCD and TmrAB are represented by cartoon and superposed.
